# Mid-Infrared Spectroscopy Study of Effects of Neonicotinoids on Forager Honey Bee (*Apis mellifera*) Fat Bodies and Their Connection to Colony Collapse Disorder

**DOI:** 10.1101/205112

**Authors:** Yuzheng Feng, Aryan Luthra, Kaiwen Ding, Yang Yang, Jordan Savage, Xinrui Wei, Roland Moeschter, Sachin Ahuja, Victor Villegas, Bogdana Torbina, Anjuli Ahooja, Tom Ellis, Tom Ellis, Anna-Maria Boechler, Andrew Roberts

**Affiliations:** Integrated Science Club, Appleby College, Oakville, ON, Canada; Canadian Light Source, University of Saskatchewan, Saskatoon, SK, Canada; University of Western Ontario, London, ON, Canada

## Abstract

This study investigated the negative effects of neonicotinoid pesticides on honey bees in environment surrounding areas of pesticide use. The aim of the experiment is to identify possible contributors to the sudden decrease in honey bee population over the past 60 years, a phenomenon known as Colony Collapse Disorder. Analysis was performed on three sets of bees: the control group which was not in contact with pesticides, the infected dead group which was a set of bees suspected to have died due to neonicotinoids, and the infected alive group which was suspected to be under the influence of neonicotinoids. After dissecting the bee samples and extracting their fat bodies, the chemical composition and protein structures of the samples were analyzed using Mid-Infrared Beamline at the Canadian Light Source. Results from the spectra of bee samples exposed to neonicotinoids demonstrated possible residual pesticide chemicals within fat bodies. Several spectral peaks were also correlated with a possible change in protein secondary structures from primarily β-sheet to α-helix within fat bodies of neonicotinoid-affected bees. It is likely that the pesticides caused the growth of additional α-helical structures, which is consistent with consequences of the inhibition of nicotinic acetylcholine receptors (nAChRs) a current pathway of harm of Colony Collapse Disorder as identified in past literature.

## 2. Introduction

Honeybee colonies are dying at an alarming rate across the world: about one-third of hives collapse every year (Dennis & Kemp, 2016). This is a pattern that has been going on in some areas for over a decade (Pic. 1). For bees and the plants that they pollinate - as well as for beekeepers, farmers, and everyone else who appreciates this marvelous social insect - this is an absolute catastrophe. The prevailing opinion suggests that Colony Collapse Disorder, hereinafter referred to as “CCD”, is linked to multi-factorial causes including pathogen infestation, beekeeping practices, and pesticide exposure in general as was identified in a 2017 CCD study (vanEngelsdorp, et al., 2017). However, a recent finding from Harvard School of Public Health (HSPH) has linked CCD with exposure to neonicotinoids, a group of systematic pesticides that appears to significantly harm honey bee colonies over the harsh winters, in which bees abandon their hives and eventually die. Although this phenomenon is quite well known, the cause is rather elusive. Many claim that it can be attributed to the multitude of causes listed above. However, many beekeepers and farmers believe that CCD should be mostly attributed to pesticides and pesticides sprayed on crops that honey bees pollinate (Aktar, Sengupta, & Chowdhury, 2009).

**Picture 1:**
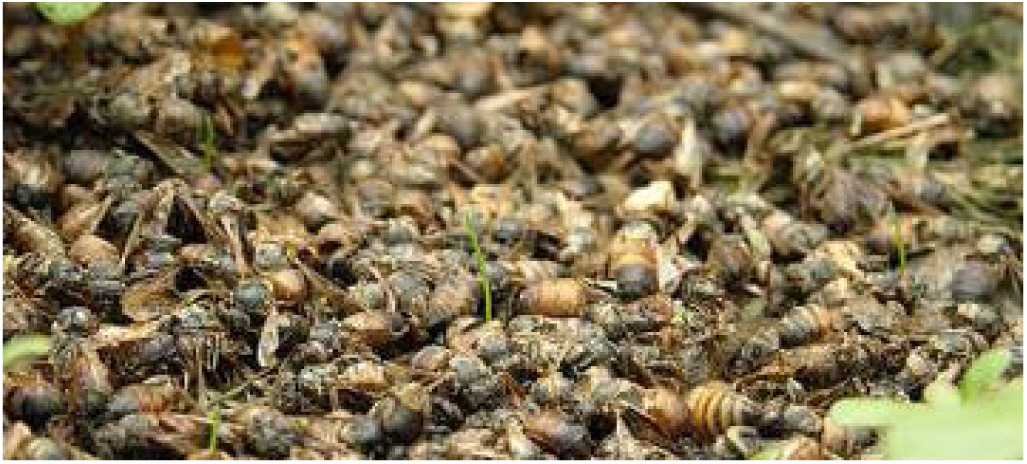
An example of a bee colony after colony collapse

The most common pesticide sprayed on crops where CCD is observed is neonicotinoids (Pic. 2). Neonicotinoids are pesticides that affect the central nervous system of insects. Many neonicotinoid pesticide products are applied to leaves and are used to treat seeds. They then accumulate in pollen and nectar of treated plants, which may be a source of exposure to pollinators. One of the first neonicotinoids that reached the market was imidacloprid, a common ingredient in garden pesticides. This product can be sprayed on the plant, but is often more effective when applied to the soil. Dinotefuran is another, more highly water-soluble, neonicotinoid that is especially good on sap-feeding insects. (Blacquière, van Gestel, & Smagghe, 2012).

**Picture 2:**
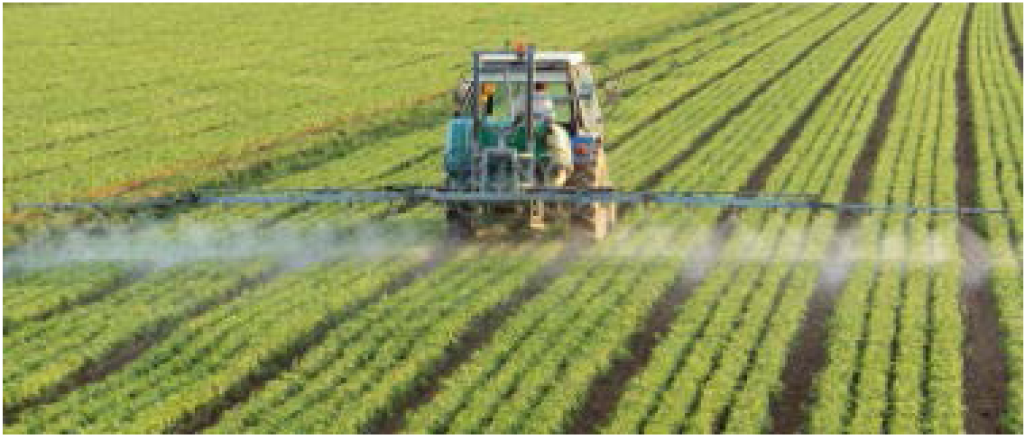
Neonicotinoids being sprayed on crops

Although it is said that these pesticides affect the nervous system, there is no understanding of which processes cause this, and how it can be prevented. This study hopes to build on these facts and contribute to information on CCD. In order to further investigate and understand the detrimental effects of neonicotinoid exposure on honey bee populations, we utilized the Mid-IR beamline at the Canadian Light Source in Saskatoon to look at the differences between the composition of the fat bodies found in contaminated bees that are at high risk of CCD and unaffected (organic) bees (through dissection). The fat body is the organ responsible for protein secretion and storage of toxins.

## 3. Theory

As it is commonly known, all light is a form of electromagnetic radiation, and visible light is a small part of this spectrum. Adjacent to the visible spectrum is UV, on the high-energy side, and Infrared (IR), on the low energy side. Mid-IR spectroscopy uses the interactions of IR light with covalent bonds to analyze composition of organic compounds. The range of the electromagnetic spectrum most useful to analyze organic compounds is the 2500-16000 nm wavelength range (Frequency: 1.9×10^13^ Hz to 1.2×10^14^ Hz). The Mid-IR Beamline allows us to use light with wavelength 1.67×10^3^ nm to 1.77×10^4^ nm.

Photons of this energy are not strong enough to ionize atoms or even excite electrons, however, they are energetic enough to induce vibrations in covalent bonds of organic compounds, this is more commonly known as “vibrational excitement.”

Once we recognize that all organic compounds undergo multiple vibrational motions (characteristic of their component atoms and respective hybridization states) and that the bonds in organic compounds act like stiff springs rather than rigid sticks or crystals, it is intuitive that all compounds will absorb IR radiation that correspond with the energy of their characteristic vibrational motions. Since the energies absorbed by a particular compound are unique to it, IR spectroscopy is a useful tool to identify the organic compounds and or functional groups present in a sample. Given enough scans and controlling all variables, one can deduce the presence, amount, sometimes even structure of organic compounds, including proteins, within the sample.

## 4. Experimental Methods

30 forager honey bees were randomly collected from each of the following environments:

– An organic environment that is pesticide-free and the bees are generally healthy. This group is labeled as the Control group.
– A region where bees are dying constantly and were thought to have a high probability of infection during their life span. Collected when dead. This group is labeled as the Infected Dead group.
– A region where bees are dying and were thought to have a high probability of infection. Collected when alive. This group is labeled as the Infected Alive group.

The bees were placed in a sealed container which was then dipped in liquid nitrogen in order to flash freeze the bees. To maintain sample integrity, the bees were then stored in a sterile, airtight, environment and stored at -45 degrees Celsius. They were not removed from storage until it was time to dissect the samples.

The dissection equipment was laid out on the table: scissors, tweezers, scoopula, and scalpel. A desiccator was placed on the table. A dissection microscope was also placed on the table. All the dissection equipment is sterilized with ethanol and Kimtech wipes. The ethanol was first squeezed onto the Kimtech wipes, and the dissection equipment was wiped with the ethanol-wetted Kimtech wipes. A frozen bee was chosen from either the Infected-dead, Infected-alive, or the control group. The bee was placed on the dissection platform. The wings and legs of the bees were cut off by small scissors in order to prevent any possible obstructions during the cutting of the thorax (anatomy shown in Fig. 1). The abdomen of the bee is then separated from the rest of the bee due to the cutting of the thorax. The end of the abdomen is then cut off by about 1-0.5mm to create an opening at the end of the abdomen. An incision was made then from the end to the front of the abdomen on the left or right side of the bee abdomen. Another incision was then made horizontally on the front of the abdomen. A tweezer was used to extend and spread the entire shell of the abdomen part into a sheet. Another tweezer was used to pick up pieces of the fat body from the contents on the sheet of abdomen shell.

**Figure 1:**
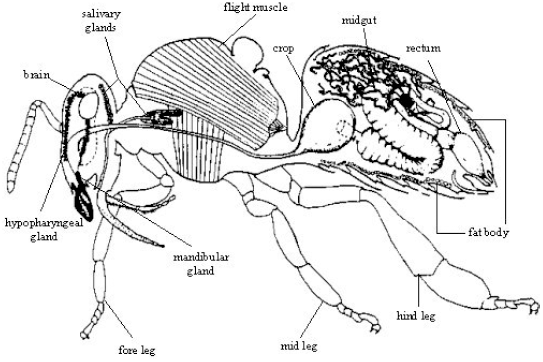
Anatomical Structure of Honey Bee

The fat body pieces were placed onto the infrared windows (BaF2). The scoopula was then used to smear the sample of fat body on the infrared window. The window was then placed on the metal window rack to be prepared for transport from the lab to the beamline. Before the transport, the sample is quickly examined under an optical microscope to ensure that it is fat body, and the thickness is appropriate. If the criteria were met (ideal specimen shown in Pic. 3), the samples were then labeled: Infected Dead (herein ID), Infected Alive (herein IA), and Control (herein C).

**Picture 3:**
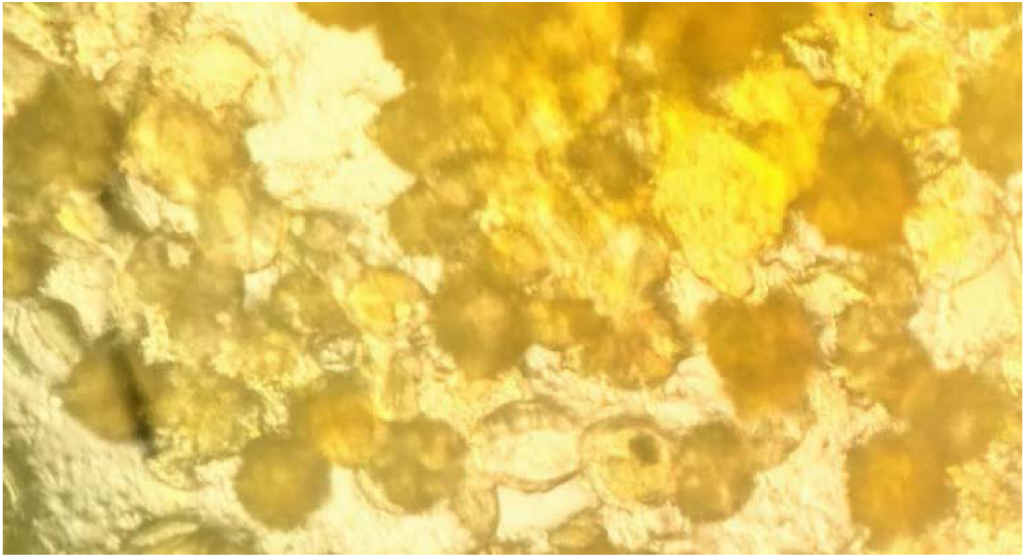
Ideal Specimen of Fat Body After Extraction

The samples are then transported to the beamlines. The slide was appropriately placed under the lens and positioned over the sample being tested. An appropriate sampling area (Diameter: 6 µm) was found on the slide using the optical lens. This area was then scanned using IR frequencies (4000-800 Wavenumbers) 255 times in order to reduce background noise. Three possible sampling areas per slide were identified.

This process was repeated for all of the sampling areas on each of the test slides. The spectra were recorded and labeled for analysis. Representative samples were chosen based on how close they were to the “average” spectra in their group. The results were analyzed.

## 5. Results

The comparison of the Mid-IR spectra of the two groups (Fig. 2) indicates a peak registered by all representative samples of the ID bees group near wavenumber 1518 cm^-1^. Control group samples, not exposed to neonicotinoids, did not reflect a significant peak on that location of the spectra. This spectra comparison (Fig. 2) also demonstrates a subtle shift in the wavenumber location of the ID samples’ Amide-I protein peak compared to the control group’s. Spectra of bee samples exposed to neonicotinoids had Amide-I peaks averaged around wavenumber 1652 cm^-1^, while control group samples registered their average peak near wavenumber 1627 cm^-1^. The Amide-I peaks of the ID samples display a smooth shape, mostly with one distinguishable peak, whereas the Amide-I regions of the Control samples contain multiple peaks of varying intensities. The spectra from wavenumber range 1400∼1800 cm^-1^ (Fig. 3) also indicates another unique peak to the ID samples near wavenumber 1378 cm^-1^.

**Figure 2:**
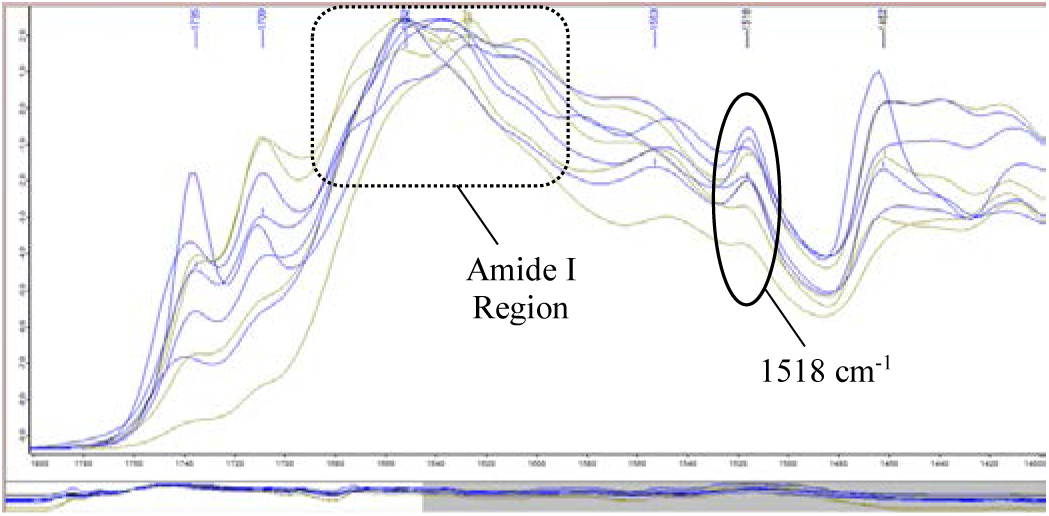
Mid-Infrared Absorbance Spectra of Infected Dead Bees Representative Samples (Blue) and Control Samples (Green) in wavenumber range 1000∼1400 cm^-1^

**Figure 3:**
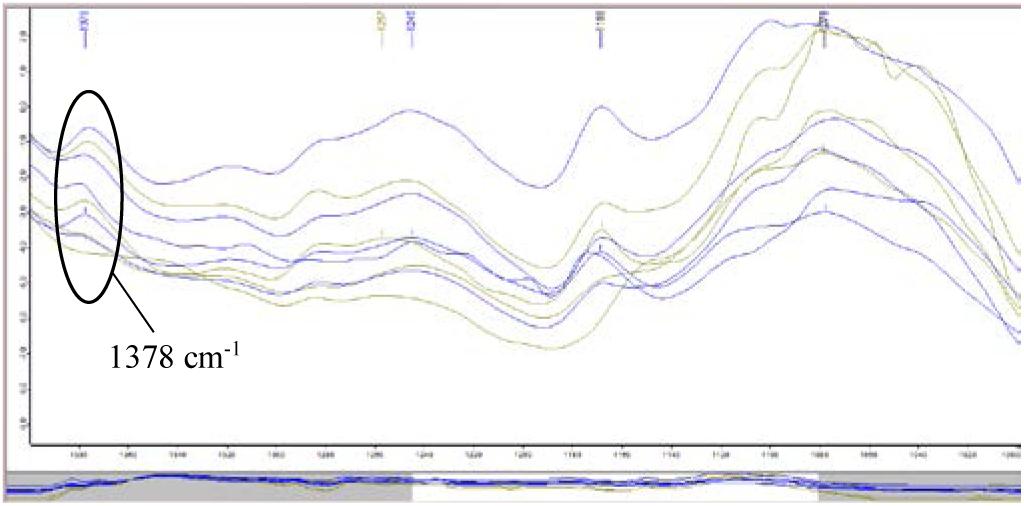
Mid-Infrared Absorbance Spectra of Infected Dead Bees Representative Samples (Blue) and Control Samples (Green) in wavenumber range 1400∼1800 cm^-1^

The exposure to neonicotinoid pesticides had a less pronounced impact on the spectra of IA samples compared to the impact on the spectra of the ID bees (Fig. 4 Fig. 5). However, 4 out of 5 of the representative samples from the IA group registered an observably intense peak near wavenumber 1515 cm^-1^. Similar smoothness near the Amide-I region as the spectra of the ID samples was indicated on the IA samples’ spectra as well (Fig. 4). Although less discernible, the spectra (Fig. 5) also demonstrates a peak in the spectra of the IA samples near wavenumber 1377 cm^-1^.

**Figure 4:**
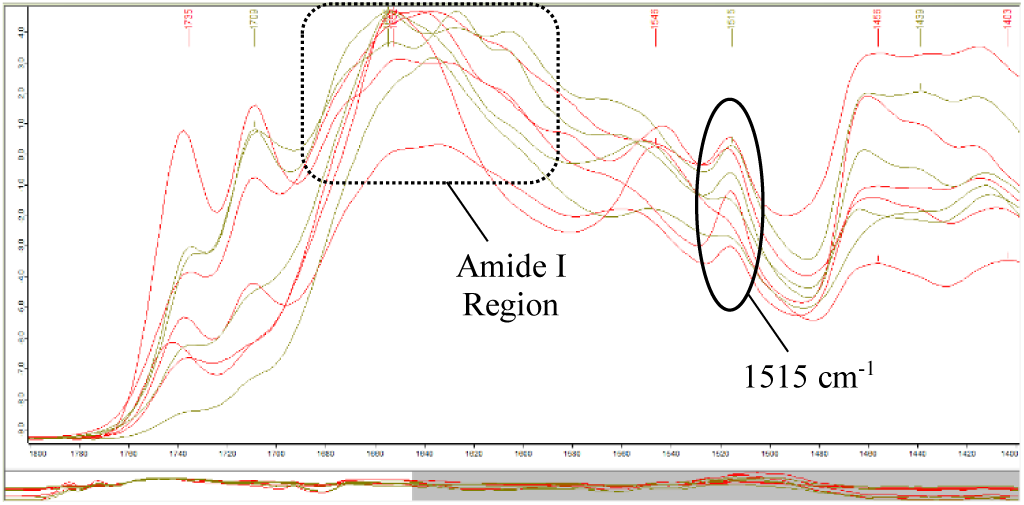
Mid-Infrared Absorbance Spectra of Infected Alive Bees Representative Samples (Red) and Control Samples (Green) in wavenumber range 1000∼1400 cm^-1^

**Figure 5:**
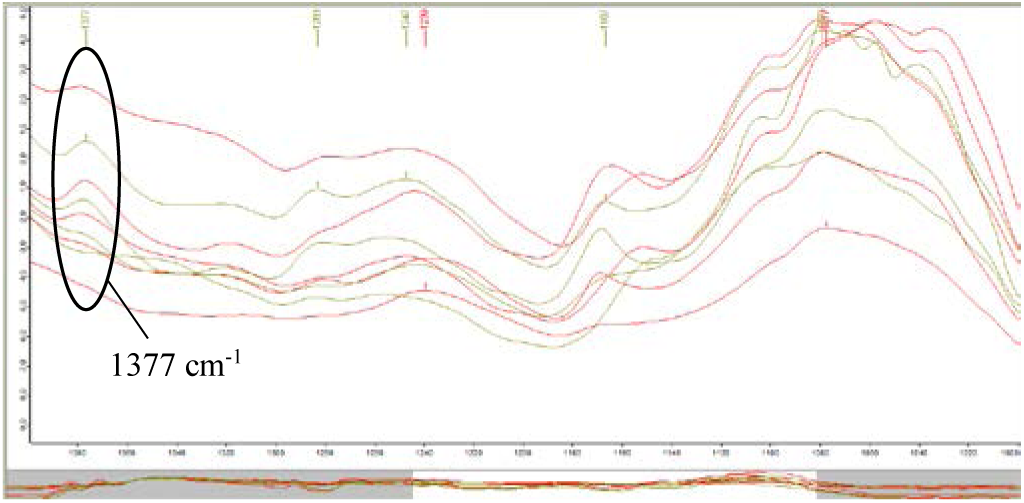
Mid-Infrared Absorbance Spectra of Infected Alive Bees Representative Samples (Red) and Control Samples (Green) in wavenumber range 1400∼1800 cm^-1^

## 6. Analysis and Discussion

From the chemical structures of the seven major types of neonicotinoids (Fig. 6), it can be identified that 4 out of the 7 types (imidacloprid, acetamiprid, nitenpyram, thiacloprid) contain the functional group 6-chloropyridine (C_5_H_3_ClN) and another 2 out of the 7 types (clothianidin, thiamethoxam) contain the function group 2-clorothiazole (C_3_H_2_ClNS). Due to the lack of pure samples of any of the above neonicotinoids, the analysis of our results was done in the aid of IR spectra of the above functional groups and of smaller constituents of these functional groups.

**Figure 6:**
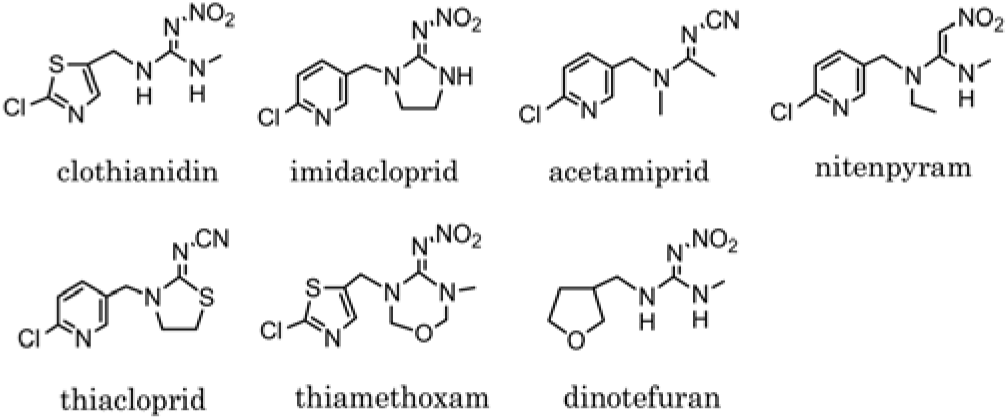
The Seven Major Types of Neonicotinoids and Their Chemical Structures (Simon-Delso et al., 2014)

In the IR spectrum of pyridine (Fig. 7), two major peaks are present at wavenumber 1579∼1590 cm^-1^ and wavenumber 1435∼1450 cm^-1^ region, respectively. A similar pattern is noted in the IR spectrum of 2-chloropyridine (Fig. 8), with two major peaks appearing at wavenumber 1572 cm^-1^ and wavenumber 1419∼1454 cm^-1^, respectively. The peaks shifted slightly to the right on the IR spectrum of 2-chloropyridine (Fig. 8) from their locations on the IR spectrum of pyridine (Fig. 7) likely due to hybridization influences from the added chlorine atom.

**Figure 7:**
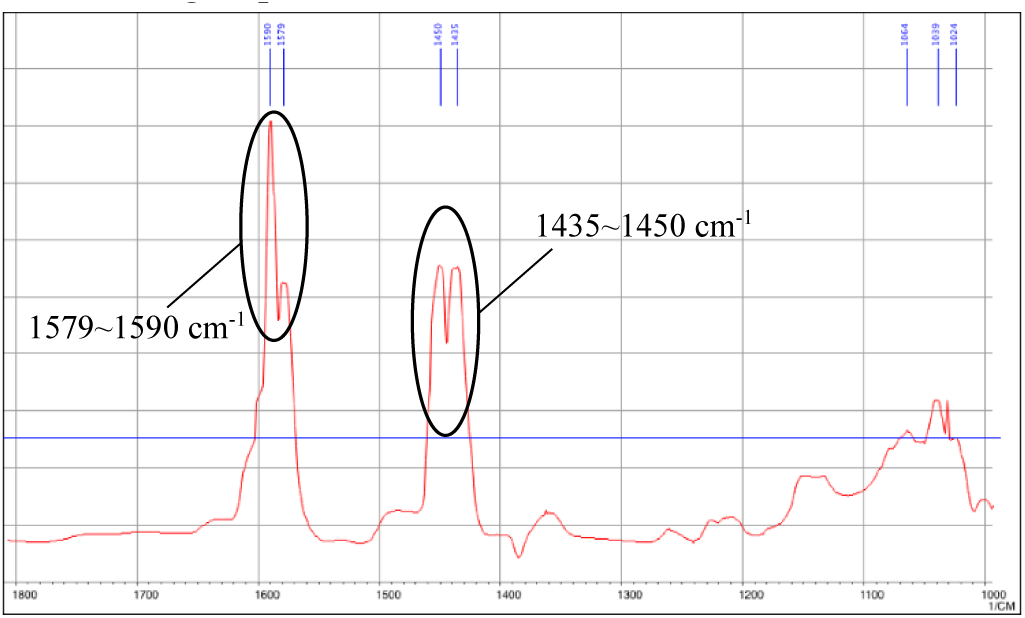
Partial IR Spectrum of Pyridine (1000∼1800 cm^-1^*) (NIST, 2016)*

**Figure 8:**
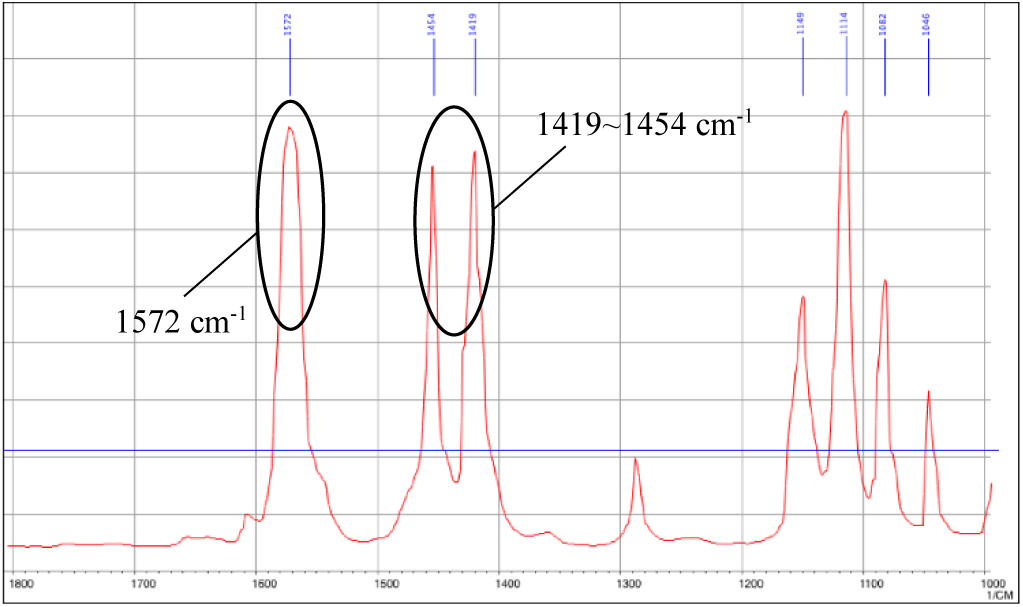
Partial IR Spectrum of 2-chloropyridine (1000∼1800 cm^-1^) (NIST, 2016)

**Table 1:**
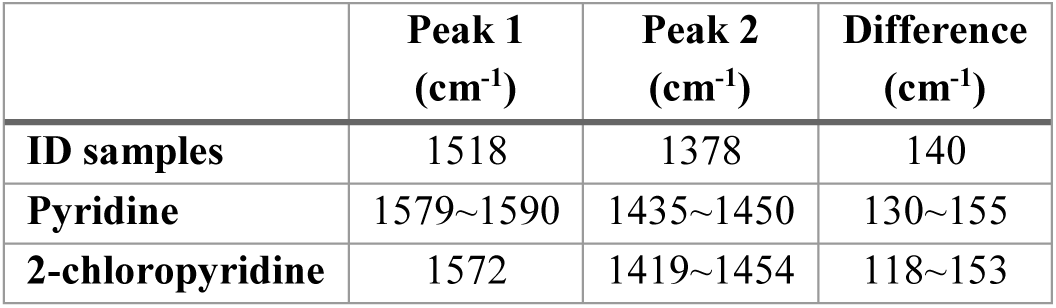
Wavenumbers of Major Peaks on IR Spectra of ID Samples, Pyridine, and 2-chloropyridine and Their Relative Differences

The two peaks identified in the results section from the IR spectra of the ID samples (Fig. 2 Fig. 3), at wavenumber 1518 cm^-1^ and 1378 cm^-1^, may not seem related at first to the above-mentioned pyridine peaks. However, further investigation was done through calculating the difference in wavenumber of the two peaks, which is 140 cm^-1^. This difference was compared to the difference in wavenumber of the two peaks identified on the IR spectrum of pyridine and 2-chloropyridine, respectively. The difference value for pyridine is 130∼155 cm^-1^, and the difference value for 2-chloropyridine is 118∼153 cm^-1^ (Tab. 1). Although not identical, these three difference values match one another, and 140 cm^-1^ falls into the range for both pyridine and 2-chloropyridine. Since the peaks identified on the IR spectra of the Infected Dead samples are also not far from the pyridine and 2-chloropyridine peaks (less than 100 cm-1), it is highly likely that the ID group’s peaks are caused by the presence of pyridine or chloropyridine in the samples. Since the main conditional difference introduced to the ID group is neonicotinoids, and 4 out of 7 types of these pesticides contain chloropyridine, the two peaks on the IR spectra of ID samples are very likely evidence of residual neonicotinoids within the fat body of ID sample bees.

A similar conclusion can be reached with peaks identified on the IR spectra of IA samples, at wavenumber 1515 cm^-1^ and 1377 cm^-1^. The difference in wavenumber of these two peaks is 138 cm^-1^, which is close to the difference value of the ID group and also fits with the difference values of the pyridine and 2- chloropyridine peaks. The IA peaks are very likely evidence of residual neonicotinoids in the system of IA sample bees.

Despite the fact that the majority of the IR absorbance of thiazole occurs in the near-IR region (below 1000 cm^-1^), a relatively significant peak was identified on the IR spectrum (Fig. 9) at wavenumber 1382 cm^-1^. This is almost an exact match to the peak location of the second peak identified on both the ID and IA spectra, at wavenumber 1378 cm^-1^ and 1377 cm^-1^ respectively. The fact that the 1382 cm^-1^ peak is relatively weak on the spectrum of thiazole may explain why the 1378 cm^-1^ and 1377 cm^-1^ peaks are not as apparent on the ID and IA spectra. Although not as convincing as the pyridine evidence, the 1378 cm^-1^ and 1377 cm^-1^ peaks are also possible evidence of the presence of thiazole in the system of sample bees. Since 2 out of the 7 types of neonicotinoids also contain chlorothiazole, these two peaks are may also be evidence of residual neonicotinoids. Neonicotinoids would only remain in the samples bees’ fat bodies if the bees failed to breakdown or release the neonicotinoids that were absorbed from the environment. The inability of the honey bees to process such chemicals point to the absence of a mechanism and possible toxicity.

**Figure 9:**
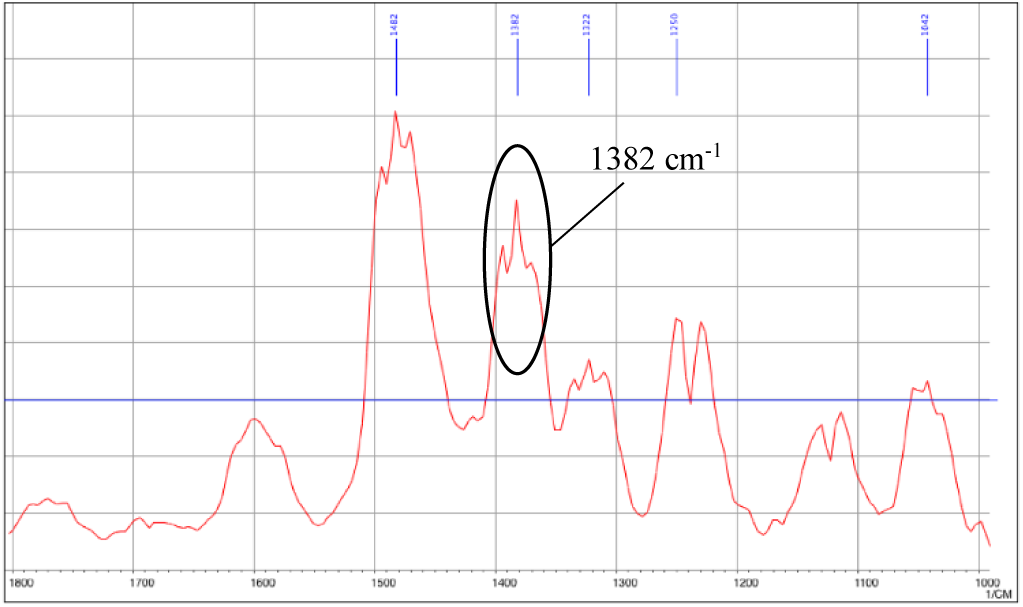
Partial IR Spectrum of Thiazole (1000∼1800 cm ^-1^) (NIST, 2016)

Although the difference in wavenumbers and similar peak locations point to the likely residual neonicotinoids within sample bees, such analysis may not be the only explanation of our results. Therefore, another possible explanation was explored to explain the ID and IA peaks.

Besides the highly likely presence of residual neonicotinoids within both ID and IA sample bees, our results also point to a possible change in the protein secondary structure of neonicotinoid-affected bees. Before discussing this change in detail, it must be explained that the analysis below was made possible by the pioneering work by Susi & Byler, who proposed a method to deconvolute the Amide-I band of protein samples into the different secondary structures using a Lorentzian shape based algorithm. Their results were verified the results with X-ray crystallography study of the proteins (Susi & Byler, 1986). In the above sample’s IR spectrum of proteins (Fig. 10), the peak shapes of different types of protein secondary structure are identified (Jabs, n.d.). Proteins with β- sheet as their most common secondary structure would register two peaks separated by approximately 60 cm^-1^ in wavenumber; however, proteins with α-helix as their secondary structure would register only one peak around wavenumber 1650∼1660 cm^-1^. In the above sample spectrum (Fig. 10), the primary secondary structure is α-helix, and the expanded band shape appears smooth with only one peak due to the influence of the intense single peak from the α-helix proteins. The presence of β-sheet and random coil is masked in this Amide I band data. Although no sample spectrum is included with β-sheet as the primary secondary structure, it is evident that the Amide I band of such a sample would have two distinct peaks due to the influence of two intense peaks caused by β-sheet proteins.

**Figure 10:**
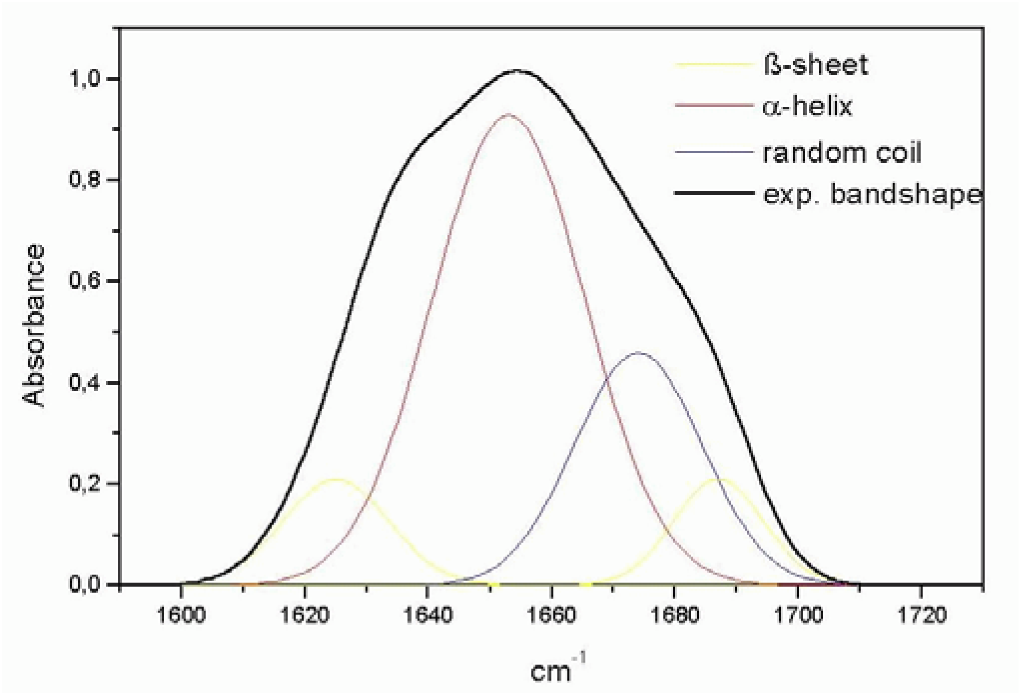
Deconvoluted IR Spectrum of Amide I Band of a Protein Sample (Jabs, n.d.)

As discussed in the results, the relatively smooth shape of the Amide I bands of the ID and IA samples’ spectra indicates that the primary secondary structure of proteins to be α-helix. The multiple peaks with varying intensities in the Control samples’ spectra point to β-sheet as the main secondary structure of proteins. This major shift in the most prevalent protein secondary structures from Control samples to neonicotinoid-affected samples indicates a probable connection to neonicotinoids. Although other factors could have influenced the shape of the Amide I bands, the relative consistency across all ID and IA samples demonstrate that a general explanation is the most likely, and a change in protein secondary structure is the best justification. Past studies by Boily et al. (2013) and van der Sluijs et al. (2013) indicate that the neonicotinoids likely cause harm through the inhibition of acetylcholine receptors within the honey bee nervous system. On the secondary level, nicotinic acetylcholine receptors (nAChRs) are primarily constituted of “two relatively conserved alpha helices within the intracellular domains of all nAChRs” (Stokes, Treinin, & Papke, 2016). The present discovery of a change in protein secondary structure in this study is consistent with this nAChRs-based pathway of harm. When neonicotinoids block nAChRs within a honey bee, the organism naturally responds by triggering growth of additional receptors, which are primarily based upon α-helices. This additional growth may be the cause behind the “refolding” see in the present study, and the “refolding” was merely caused by additional proteins instead of changes to existing ones. However, the nature of the study as an IR-based experiment made it impossible for to identify the exact types of proteins being influenced by this change, but further experiments must be performed using X-ray crystallography to investigate possible connections to nAChRs.

Due to the presently unclear connection to acetylcholine receptors, this study proposes an alternate explanation for the observed change from β-sheet to α-helix. The aggregation of α- helical structures has been associated with the membrane-mediated growth of amyloid fibrils. This growth is the hallmark of a class of human diseases known as amyloidosis, which includes Alzheimer’s disease, Parkinson’s disease, and type II diabetes mellitus. These diseases are highly debilitating and associated with misfolding on the secondary level of protein structures. Although an aggregation of β-sheet fibrils has been identified as the indicator for amyloidosis development, an α- helical structure has been found to catalyze protein misfolding in the onset of amyloidosis. In their 2008 study, Apostolidou, Jayasinghe, and Langen investigated the misfolding of the human islet amyloid polypeptide (hIAPP), the primary amyloidogenic agent in type II diabetes mellitus. Via Electron Paramagnetic Resonance (EPR) spectroscopy, it was found that “the formation of α-helical, membrane-bound hIAPP exposes a highly amyloidogenic stratch of hIAPP to an environment that is expected to promote the formation of misfolded β-sheet structures” (Apostoulidou, Jayasinghe, & Langen, 2008). Beyond this promotion of β-sheet fibrils growth, the 2008 study also references another research in the journal *Biochemistry*, which suggested that α-helical, membrane-bound hIAPP has the capacity to oligomerize, and such oligomers could then transition to misfolded β-sheet fibrils (Knight, Hebda, & Miranker, 2006). This finding is further explored by Miranker and Magzoub in 2012 using fluorescence correlation spectroscopy (FCS). They identified that even before the formation of misfolded β-sheet structures, the α-helical mediated oligomerization allows the membrane-bound hIAPP to “acquire cell-penetrating peptide (CPP) properties, facilitating access to the mitochondrial compartment, resulting in its dysfunction” (Magzoub & Miranker, 2012). In the three studies above, it can be seen that understanding of the role of α- helical structures in the debilitating nature of amyloidosis has advanced an evolved. In early-stage amyloidosis onset, aggregation of α-helical proteins seems crucial in the facilitation of β-sheet misfolding. However, as presented by Magzoub and Miranker, cellular damage begins before the growth of β-sheet structures. Their result is also corroborated by another study employing computation biophysics, also identifying the permeabilization of the membranes as the pathway of toxicity by hIAPP (Pannuzzo, et al., 2013). The transition from β-sheet to α-helix as the prevalent secondary structure – identified in this present research – may be caused by an aggregation of hIAPP-like α-helical membrane-bound polypeptides, which could cause membrane permeabilization, exposing cellular contents to external damage. This exposure and its subsequent toxicity does not necessitate the growth of β- sheet fibrils, which may explain the absence of excessive β- sheet growth.

Various recent studies also identify neonicotinoids as the probable culprit behind colony collapse disorder, including the Harvard School of Public Health study by Lu, Warchol & Callahan, and an 18-year distribution study by Woodcock, et al. in England. The Lu study identified impairment of winterization as a major pathway through which neonicotinoids lead to colony collapse disorder (Lu, Warchol, & Callahan, 2014). The British study did not identify any particular pathway of colony collapse disorder; instead, it related distribution of wild bees to that of amounts of neonicotinoid use in oilseed rape and identified the long-term negative impacts of these pesticides on various species of bees (Woodcock, et al., 2016). The results of this study are in agreement with both studies by Lu and Woodcock’s teams. This experiment not only indicates the likely presence of residual neonicotinoids in ID and IA samples’ fat bodies, it also generates to a novel hypothesis that colony collapse disorder could be the result additional growth of α-helical structures – leading to an observed shift in the most prevalent protein secondary structure.

## 7. Conclusions

By comparing spectra of the ID and IA groups to the spectra obtained from literature on 2-chloropyrdine and thiazole, it can be said that neonicotinoids are likely present in the fat bodies of the treated bees. However, they were not present in the control samples, indicating an accumulation of the pesticide within fat bodies of the affected bees. Although this conclusion is highly likely, it remains as one possible explanation of the observed spectral peak differences.

The shape of the Amide-1 peaks in sample spectra and their differences across treatments indicate that there has likely been a shift in the most prevalent protein secondary structures in the fat bodies of the ID and IA groups. This discovery of a change in protein secondary structure from β-sheet to α-helix could be consistent with the nAChRs pathway –proposed by past research – by which neonicotinoids induce harm to honey bees. This shift in observed secondary structures could also be caused by the growth of other α-helical structures, potentially including amyloidogenic membrane-bound polypeptides.

## 8. Further Research

Further experiments must be done to confirm our results and possibly verify our hypothesis. First, a more controlled experiment would be performed, with controlled amount of sample bees exposed and the types and concentrations of neonicotinoids the samples are exposed to. This would allow us to perform qualitative analysis of the sample bees’ behaviour and also have a better understanding of the peaks we would be expecting from the IR spectra. Control over the type of neonicotinoids would also make it possible to scan the exact pure sample of the pesticide so that its IR spectrum may be used to compare with result spectra.

Second, proper X-ray crystallography studies would be ideal. It will allow for the identification of the exact proteins involved in the secondary structure change, in order to investigate possibilities of the two scenarios presented above (nAChRs inhibition or amyloidosis onset). Although the details of further research are yet to be determined, these studies will certainly help illustrate a more comprehensive picture of neonicotinoids’ relationship to colony collapse disorder.

## References

1. Aktar, M. W., Sengupta, D., &Chowdhury, A. (2009). Impact of pesticides use in agriculture: their benefits and hazards. Interdisciplinary Toxicology, 2(1), 1–12. http://doi.org/10.2478/v10102-009-0001-7

2. Apostolidou, M., Jayasinghe, S. A., &Langen, R. (2008). Structure of a-Helical Membrane-bund Human Islet Amyloid Polypeptide and Its Implications for Membrane-mediated Misfolding. Journal of Biological Chemistry, 283(25), 17205–17210. doi:10.1074/jbc.M801383200

3. Blacquière, T., Smagghe, G., van Gestel, C.A.M. et al. (2012) Neonicotinoids in bees: a review on concentrations, side-effects and risk assessment. Ecotoxicology, 21(973), 973–992. https://doi.org/10.1007/s10646-012-0863-x

4. Boily, M., Sarrasin, B., DeBlois, C., Aras, P., &Chagnon, M. (2013). Acetylcholinesterase in honey bees (*Apis mellifera*) exposed to neonicotinoids, atrazine and glyphosate: laboratory and field experiments. Environmental Science and Pollution Research, 20(8), 5603–5614.

5. Byler, D. M., &Susi, H. (1986). Examination of the Secondary Structure of Proteins by Deconvolved FTIR Spectra. Biopolymers, 25, 469–487.

6. Dennis, B., Kemp, W.P. (2016) How Hives Collapse: Allee Effects, Ecological Resilience, and the Honey Bee. PLoS ONE 11(2): e0150055. https://doi.org/10.1371/journal.pone.0150055

7. Jabs, A. (n.d.). Determination of Secondary Structure in Proteins by Fourier Transform Infrared Spectroscopy (FTIR). Retrieved December 30, 2016, from Jena Library of Biological Macromolecules: http://jenalib.leibniz-fli.de/ImgLibDoc/ftir/IMAGE_FTIR.html

8. Knight, J. D., Hebda, J. A., &Miranker, A. D. (2006). Conserved and cooperative assembly of membrane-bound alpha-helical states of islet amyloid polypeptide. Biochemistry, 45(31), 9496–9508. doi:10.1021/bi060579z

9. Magzoub, M., &Miranker, A. D. (2012). Concentration-dependent transitions govern the subcellulr localization of islet amyloid polypeptide. The FASEB Journal, 26(3), 1228–1238. doi:10.1096/fj.11-194613

10. Lu, C., Warchol, K. M., &Callahan, R. A. (2014). Sub-lethal exposure to neonicotinoids impaired honey bees winterization before proceeding to colony collapse disorder. Bulletin of Insectology, 1(1), 125–130.

11. Pannuzzo, M., Raudino, A., Milardi, D., Rosa, C. L., &Karttuen, M. (2013). a-Helical Structures Drive Early Stages of Self-Assembly of Amyloidogenic Amyloid Polypeptide Aggregate Formation in Membranes. Scientific Reports, 3(2781). doi:10.1038/srep02781

12. Stokes, C., Treinin, M., &Papke, R. L. (2015). Looking below the surface of nicotinic actylcholine receptors. Trends in Pharmocological Sciences, 8(8), 514–523. doi:10.1016/j.tips.2015.05.002

13. van der Sluijs, J. P., Simon-Delso, N., Goulson, D., Maxim, L., Bonmatin, J.-M., &Belzunces, L. P. (2013). Neonicotinoids, bee disorders and the sustainability of pollinator services. Current Opinion in Environmental Sustainability, 5(3-4), 293–305.

14. vanEngelsdorp, D., Traynor, K.S., Andree, M., Lichtenberg, E.M., Chen, Y., Saegerman, C., et al. (2017) Colony Collapse Disorder (CCD) and bee age impact honey bee pathophysiology. PLoS ONE 12(7): e0179535. https://doi.org/10.1371/journal.pone.0179535

15. Woodcock, B.A., Isaac, N. J., Bullock, J. M., Roy, D. B., Garthwaite, D. G., Crowe, A., &Pywell, R. F. (2016). Impacts of neonicotinoid use on long-term population changes in wild bees in England. Nature Communications.

